# Increased migratory/activated CD8+ T cell and low avidity SARS-CoV-2 reactive cellular response in post-acute COVID-19 syndrome

**DOI:** 10.1101/2022.12.03.519007

**Authors:** Krystallenia Paniskaki, Margarethe J. Konik, Moritz Anft, Harald Heidecke, Toni L. Meister, Stephanie Pfaender, Adalbert Krawczyk, Markus Zettler, Jasmin Jäger, Anja Gaeckler, Sebastian Dolff, Timm H. Westhoff, Hana Rohn, Ulrik Stervbo, Carmen Scheibenbogen, Oliver Witzke, Nina Babel

**Author notes:** Equal contribution (Krystallenia Paniskaki, Margarethe J. Konik). Correspondence should be addressed to: Dr Krystallenia Paniskaki.

## Abstract

The role of autoimmunity in post-acute sequelae of COVID-19 (PASC) is not well explored, although clinicians observe a growing population of convalescent COVID-19 patients with manifestation of post-acute sequelae of COVID-19. We analyzed the immune response in 40 post-acute sequelae of COVID-19 patients with non-specific PASC manifestation and 15 COVID-19 convalescent healthy donors. The phenotyping of lymphocytes showed a significantly higher number of CD8+ T cells expressing the Epstein-Barr virus induced G protein coupled receptor 2, chemokine receptor CXCR3 and C-C chemokine receptor type 5 playing an important role in inflammation and migration in PASC patients compared to controls. Additionally, a stronger, SARS-CoV-2 reactive CD8+ T cell response, characterized by IFNγ production and predominant T_EMRA_ phenotype but low SARS-CoV-2 avidity was detected in PASC patients compared to controls. Furthermore, higher titers of several autoantibodies were detected among PASC patients. Our data suggest that a persistent inflammatory response triggered by SARS-CoV-2 might be responsible for the observed sequelae in PASC patients. These results may have implications on future therapeutic strategies.

## Introduction

Post-infectious Myalgic Encephalomyelitis/Chronic Fatigue Syndrome (ME/CFS) is already described as a sequelae after numerous primarily viral infections (Epstein-Barr virus, cytomegalovirus, etc.) and SARS-CoV-2 virus is lately adding to the list (1). Post-acute sequelae of COVID-19 presents as a chronic multisystemic disease characterized by a variety of respiratory, cardiovascular, gastroenterological and neurological symptoms (2–5). A growing body of literature suggests that a combination of virus and host factors, residual inflammation (1,6–7), microvascular dysregulation/endothelial injury (8–10), autoimmune phenomena and abnormal cellular energy metabolism (11–12) contribute to PASC.

Emerging epidemiologic studies (13–14) draw attention to concentration disorders, posttraumatic stress disorder, sleep disorders as psychiatric/neurological sequelae after COVID-19 disease. To this end, several studies demonstrated anatomical changes in structures of the brain of COVID-19 survivors with post-acute COVID-19 (15–17).

Similar to ME/CFS immune alterations have been also demonstrated to be associated with PASC. Thus, distinct immune changes have newly been described in PASC with primarily lung sequelae. Cheon et al showed that functional SARS-CoV-2-specific memory T and B cells are abundant in bronchial lavage fluid of COVID-19 adults with primarily lung sequelae compared to those of blood (18). Vijayakumar et al demonstrate distinct immune and proteomic changes in post-COVID-19 airways, where increased bronchoalveolar lavage cytotoxic T cells are linked to epithelial damage and airway disease (19). Few studies approach experimentally the immune mechanisms responsible for system specific PASC separately-gastroenterological, respiratory, cardiovascular or neurological PASC. However, data on the immune pathogenesis of PASC as a disease entity (non-specific PASC), which affects a significant proportion of the general population are currently missing. To address these knowledge gaps and the contribution of immunity, including humoral and cellular response in non-specific PASC pathogenesis, we performed a comprehensive immune profiling of 40 patients with non-specific PASC. As control we used 15 healthy COVID-19 convalescent adults.

## Materials and methods

### Study participants

We used peripheral blood mononuclear cells (PBMCs) and serological samples from 40 convalescent COVID-19 patients with post COVID-19 syndrome (further referred as PASC) and 15 convalescent COVID-19 patients without clinical manifestation of post COVID-19 syndrome (further referred as control). The clinical criteria of post-acute COVID-19 syndrome as defined by Nalbandian et al (2) and NICE guidelines (https://www.nice.org.uk/guidance/ng188/resources/covid19-rapid-guideline-managing-the-longterm-effects-of-covid19-pdf-51035515742) were applied to set the diagnosis of post COVID-19 syndrome and therefore recruitment of the study participants. Cognitive/psychiatric impairment/symptomatic was based on ICD 10 psychiatric relevant diagnoses. Demographic and clinical characteristics are provided in tables 1–2.

**Table 1:** Clinical Symptoms and comorbidities of the study cohorts

|  |  | PASC N(%) | Control N(%) |
| --- | --- | --- | --- |
| **PASC Type** | Ongoing COVID-19 | 0 (0) | N/A |
|  | Post-acute COVID-19 Syndrome | 40 (100) | N/A |
| **Post-acute COVID-19 Symptoms** | Shortness of breath/exercise intolerance | 28 (70) | N/A |
|  | Cognitive disturbances (brain fog) | 23 (57.5) | N/A |
|  | Fatigue | 17 (42.5) | N/A |
|  | Decline of life quality | 16 (40) | N/A |
|  | Dyspnea | 11 (27.5) | N/A |
|  | Muscular weakness | 9 (22.5) | N/A |
|  | Anxiety/depression | 7 (17.5) | N/A |
|  | Sleep disturbances | 6 (15) | N/A |
|  | Chest pain | 4 (10) | N/A |
|  | Joint pain | 3 (7.5) | N/A |
|  | Cough | 3 (7.5) | N/A |
|  | Headaches | 3 (7.5) | N/A |
|  | Smell/taste disturbance | 3 (7.5) | N/A |
|  | Posttraumatic stress disorder | 2 (5) | N/A |
|  | Thromboembolism | 2 (5) | N/A |
|  | Persistent oxygen requirement | 0 (0) | N/A |
|  | Palpitations | 0 (0) | N/A |
|  | Hair loss | 0 (0) | N/A |
| **Comorbidities prior to COVID-19** | Obesity | 33 (82.5) | 3 (20) |
|  | Arterial hypertension | 9 (22.5) | 0 (0) |
|  | Asthma bronchiale | 4 (10) | 0 (0) |
|  | Malignity | 3 (7.5) | 0 (0) |
|  | Autoimmune disease | 3 (7.5) | 0 (0) |
|  | Depression | 3 (7.5) | 0 (0) |
|  | Diabetes | 3 (7.5) | 0 (0) |
|  | Coronary heart disease | 1 (2.5) | 0 (0) |

**Table 2:** Demographic and clinical characteristics of the study cohorts

|  | **PASC (N=40)** | **Control (N=15)** |
| --- | --- | --- |
| Age years -median (range) | 51.5 (19-68) | 30 (19-50) |
| Female gender N (%) | 25 (63) | 10 (70) |
| BMI -median (range) | 29 (17.6-52.5) | 22 (17.9-35) |
| Time since COVID-19 Diagnosis (months) | 10 (2-16) | 8 (2-12) |
| Duration of PASC symptoms (months)* | 10 (2-16) | 0 (0-0) |
| COVID-19 Severity N (%) |  |  |
| *Moderate* | 38 (96) | 15 (100) |
| *Severe* | 1 (2) | 0 (0-0) |
| *Critical* | 1 (2) | 0 (0-0) |
| Time last vaccination up to recruitment (months)** | 4 (1-8) | 4 (1-8) |
| Variant of concern N (%) |  |  |
| *WT* | 25 (63) | 8 (53) |
| *Alpha* | 13 (32) | 1 (7) |
| *Delta* | 1 (2.5) | 4 (27) |
| *Omicron* | 1 (2.5) | 2 (13) |

\*Months from SARS-CoV-2 Diagnosis (positive PCR) till the time point of study recruitment
\*\*Months from last vaccination up to study recruitment

### Preparation of PBMCs

As previously described, peripheral blood was collected in S-Monovette K3 EDTA blood collection tubes (Sarstedt)(20–21). Collected blood was prediluted in PBS/BSA (Gibco) at a 1:1 ratio and underlaid with 15 mL of Ficoll-Paque Plus (GE Healthcare). Tubes were centrifuged at 800g for 20 min at room temperature. Isolated PBMCs were washed twice with PBS/BSA and stored at −80 °C until use. The cryopreserved PBMCs were thawed by incubating cryovials 2-3 minutes at 37 °C in bead bath, washed twice in 37°C RPMI 1640 media (Life Technologies) supplemented with 1% penicillin-streptomycin-glutamine (Sigma-Aldrich), and 10% fetal calf serum (PAN-Biotech) medium, and incubated overnight at 37 °C.

### Flow cytometry-Characterization T cell activation/migration status

Thawed and rested overnight PBMCs were plated in 96-U-Well plates in RPMI 1640 media (Life Technologies). The PBMCs were stained with optimal concentrations of antibodies for 10 min at room temperature in the dark. Stained cells were washed with PBS/BSA and were immediately acquired on a CytoFLEX flow cytometer (Beckman Coulter) (gating strategy fig. S1). Fluorescence minus one controls were used for optimal gating of Epstein-Barr virus induced G protein coupled receptor 2 (EBI2), chemokine receptor CXCR3 (CXCR3) and C-C chemokine receptor type 5 (CCR5) populations. No modification to the compensation matrices was required throughout the study. Detailed listing of the antibody panel for the characterization of EBI2+, CCR5+ and CXCR3 T cells is presented in supplementary table S1.

### Flow cytometry - Measurement of SARS-CoV-2 reactive T cells

As previously described, PBMCs were plated in 96-U-Well plates in RPMI 1640 media (Life Technologies) (20)(21). Each well was stimulated with one of the following SARS-CoV-2 proteins: the complete sequence of B.1.1.7 D614G Spike mutant (alpha) (JPT Peptide Technologies) or wildtype (WT) S-protein (Miltenyi Biotec) or left untreated as a control for 16 h. The proteins were dissolved per manufacturer’s directions. As a positive control, cells were stimulated with staphylococcal enterotoxin B (1 μg/mL, Sigma-Aldrich). After 2 h, brefeldin A (1 μg/mL, Sigma-Aldrich) was added. Detailed listing of the antibody panels for general phenotyping and T cell activation *ex vivo* is in Table S2. The PBMCs stimulated overnight were stained with optimal concentrations of antibodies for 10 min at room temperature in the dark. Stained cells were washed twice with PBS/BSA before preparation for intracellular staining using the Intracellular Fixation & Permeabilization Buffer Set (Thermo Fisher Scientific) as per the manufacturer’s instructions. Fixed and permeabilized cells were stained for 30 min at room temperature in the dark with an optimal dilution of antibodies against the intracellular antigen. All samples were immediately acquired on a CytoFLEX flow cytometer (Beckman Coulter) (gating strategy fig. S2). Quality control was performed daily using the recommended CytoFLEX daily QC fluorospheres (Beckman Coulter). No modification to the compensation matrices was required throughout the study. Antigen-reactive responses were considered positive after the non-reactive background was subtracted, and more than 0.01% were detectable. Negative values were set to zero.

### SARS-CoV-2 neutralization assay

As previously described (21), for the virus neutralization assay, sera were incubated for 30 min at 56°C in order to inactivate complement factors. Single cycle VSV*ΔG(FLuc) pseudoviruses bearing the SARS-CoV-2 WT Spike (D614G) protein or SARS-CoV-2 B.1.1.7. (alpha) Spike protein in the envelope were incubated with quadruplicates of two-fold serial dilutions of immune sera in 96-well plates prior to infection of Vero E6 cells (1×10^4^ cells / well) in DMEM + 10% FBS (Life Technologies). At 18 hours post infection, firefly luciferase (FLuc) reporter activity was determined using a CentroXS LB960 (Berthold) and the reciprocal antibody dilution causing 50% inhibition of the luciferase reporter was calculated (PVND50).

### Autoantibodies

Human IgG antibodies (Ab) against 12 different G protein-coupled receptors; beta-2 adrenergic receptor (ß2-adr-R), bradykinin receptor B1 (Brady-R1), angiotensin II receptor type 1 (ATR1), angiotensin II receptor type 2 (ATR2), Mas 1 receptor (MasR), adrenoceptor alpha 1A (a1-adr-R), CXCR3, endothelin A receptor (ETAR), endothelin B receptor (ETBR), M5 muscarinic acetylcholine receptor (M5R), protease activated receptor 1 (PAR1), protease activated receptor 2 (PAR2), two proteins; angiotensin-converting enzyme 2 (ACE2), Annexin 2 and the transmembrane receptors Annexin 2R and stabilin 1 (STAB1) were detected from frozen serum using commercial ELISA kits (CellTrend, Luckenwalde, Germany) according to the manufacturer’s instructions as previously described(22).

### Statistics

Flow cytometry data were analyzed using FlowJo version 10.6.2 (BD Biosciences); gating strategies are presented in figure S1 and S2. For the analysis of anti-SARS-CoV-2 reactive T cells, a threshold of 0.01% was employed to define a detectable response. Single stains and fluorescence-minus-one controls were used for gating. Gates of each study participant were adjusted according to the negative control. CD4+ T cells expressing CD154 and CD137 and CD8+ T cells expressing CD137 were defined as reactive T cells. Statistical analysis was performed using GraphPad Prism v7. Categorical variables are summarized as numbers and frequencies; quantitative variables are reported as median and interquartile range. Normality Tests were performed with Shapiro-Wilk test. All applied statistical tests are two-sided. Frequencies of SARS-CoV-2-protein reactive T cells in the PASC study group and the control group were compared using exact two-tailed Mann-Whitney test. The age between the two cohorts was compared using unpaired two-tailed t-test, and gender was compared using two-tailed Fisher’s exact test. p values below 0.05 were considered significant; only significant p values are reported in the figures. p values were not corrected for multiple testing, as this study was of an exploratory nature.

## Results

### 1. Characterization of the study groups

Our study group comprised 40 convalescent COVID-19 patients with non-specific PASC (further referred as PASC) and 15 convalescent COVID-19 patients without clinical manifestation of post COVID-19 syndrome (further referred as control). The clinical criteria of PASC as defined by Nalbandian et al (2) and NICE guidelines were applied to set the diagnosis of post COVID-19 syndrome and therefore recruitment of the study participants. All study participants had a negative SARS-CoV-2 nasal swab tested via PCR on recruitment. During the acute phase of COVID-19 disease, 100% (n=15) and 96% (n=38) of the control and PASC group respectively, presented moderate COVID-19 disease severity without need for hospitalization, whereas only 4% (n=2) were severely or critically ill and hospitalized. The clinical symptoms of PASC are summarized in table 1.

The median COVID-19 convalescence time for the control and PASC study groups at the timepoint of the recruitment was 8 (range 2-12) and 10 (range 2-16) months respectively. All control individuals (n=15) and 82.5% (n=33) of the PASC study group received at least two COVID-19 mRNA vaccinations (either prior or after the infection). The latest COVID-19 vaccination took place in a median time of 4 (range 1-8) months before the study recruitment for both study groups.12.5% (n=7) of the PASC study group were not vaccinated against SARS-CoV-2 neither before nor after the infection. The majority of the PASC patients and control study participants were infected with either the WT or the alpha variant (table 2).

The median age of the PASC study group was 51.5 years (range 19-68 years), whereas the control cohort was significantly younger, with a median age of 30 years (range 19-50 years, p=0.0002 two tailed unpaired t test). The PASC and control cohorts comprised of 63% (n=25) and 70% (n=10) female participants, respectively and showed no statistical gender difference (Fisher’s exact test, p<0.05). PASC patients (median BMI 29, range 17.6-52.5) showed significantly higher BMI compared to controls (median BMI 22, range 17.9-35) (Mann Whitney test, p=0.0003). All PASC study subjects suffered from at least two symptoms. The PASC study group presented significantly more comorbidities compared to the controls (unpaired t test p=0.0006). The demographic and clinical characteristics of the study cohorts are presented in Tables 1–2.

### 2. Increased frequencies of EBI2-, CXCR3-, and CCR5-expressing T cells in PASC patients suggests a high inflammatory and migratory T cell response

Firstly, we characterized the migration/activation status of the peripheral T cells. We detected similar frequencies of CD4+ and CD8+ T cells among the two study groups (fig. 1A & 1E). In order to assess the activation und migration status of the peripheral T cells, we analyzed three G protein-coupled receptors crucial for inflammation and autoimmunity: the chemokine receptors CCR5 and CXCR3, and the EBI2.

**Figure 1:**
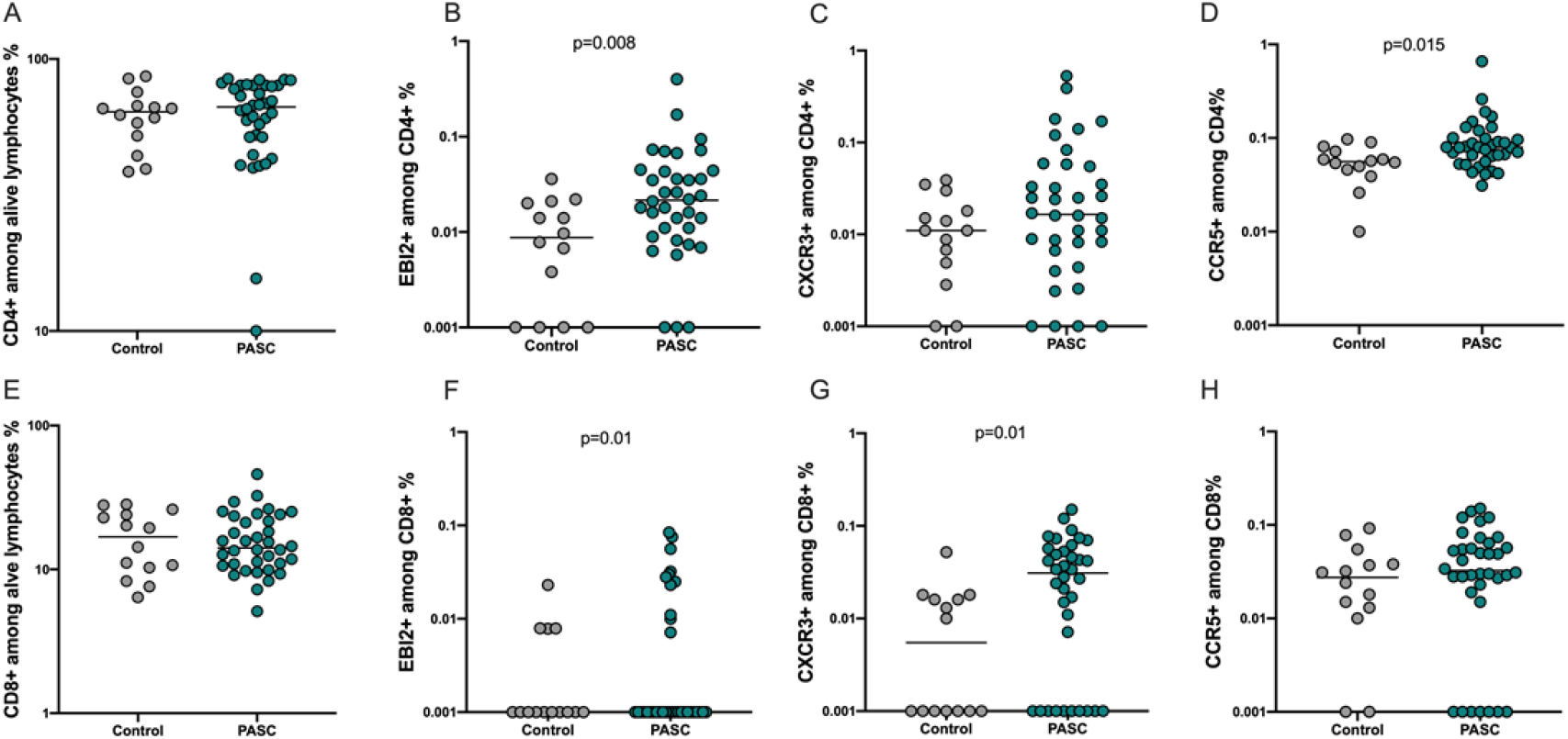
CD4+ and CD8+ T cells with a high inflammatory and migratory profile among the PASC patients. The expression of the activation and migration markers EBI2, CCR5 and CXCR3 was assessed on CD4+ and CD8+ T cells of 40 PASC patients and 15 convalescent controls. (A) CD4+ T cell frequencies of PASC subjects compared to controls. (B-D) Frequencies of CD4+EBI2+, CD4+CXCR3+ and CD4+CCR5+ T cells of PASC individuals compared to controls. (E) CD8+ T cell frequencies of PASC subjects compared to controls. (F-H) Frequencies of CD8+EBI2+, CD8+CXCR3+ and CD8+CCR5+ T cells of PASC individuals compared to controls. Scatterplots show line at median. Unpaired data were compared with Mann-Whitney-test. P<0.05 was considered significant, only significant p values are documented in the figures.

The PASC study group showed significantly higher frequencies of CD4+ and CD8+ T cells expressing EBI2, CD4+ T cells expressing CCR5, CD8+ T cells expressing CXCR3, and higher CD4+CXCR3+ T cell frequencies without reaching a statistical significance though compared to control group (fig. 1B-D & 1F-G). CD8+CCR5+ T cells showed similar frequencies among PASC and control study group (fig. 1H). We also observed a significantly higher frequencies of CD4+ and CD8+ T cells co-expressing CCR5 and CXCR3 molecules, and CD4+ T cells co-expressing CCR5 and EBI2 molecules in PASC patients compared to controls(fig. 2).

**Figure 2:**
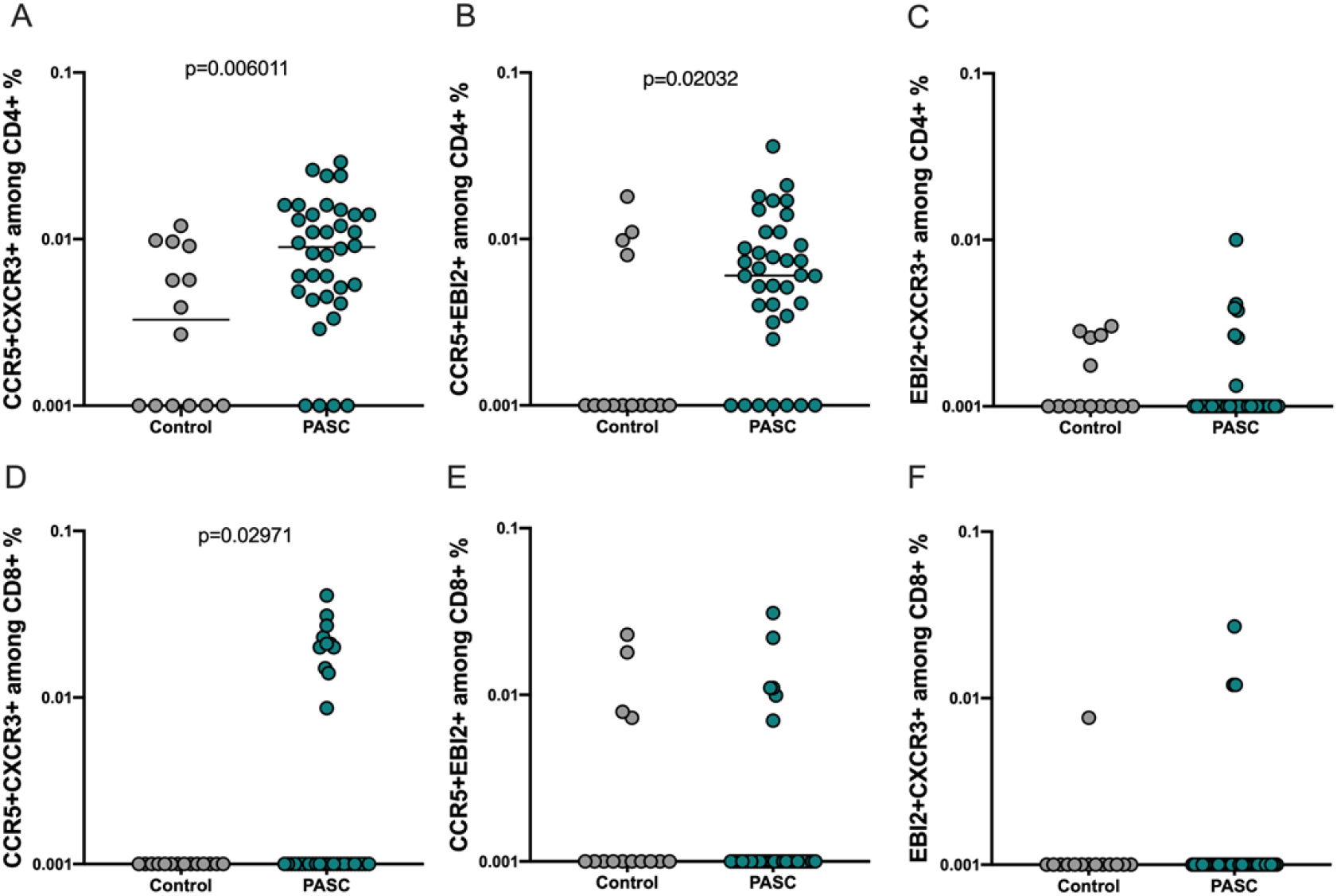
Increased frequencies of double positive CCR5+CXCR3+CD4+ and CCR5+CXCR3+CD8+ T cells among PASC patients. Analysis of the simultaneous expression of EBI2, CCR5 or/and CXCR3 receptor of T cell surface. (A-C) Frequencies of CD4+CCR5+CXCR3+, CD4+CCR5+EBI2+ and CD4+EBI2+CXCR3+ T cells among PASC and controls. (D-F) CD8+CCR5+CXCR3+, CD8+CCR5+EBI2+ and CD8+EBI2+CXCR3+ T cells among PASC and controls. Scatterplots show line at median. Unpaired data were compared with Mann-Whitney-test. P<0.05 was considered significant, only significant p values are documented in the figures.

### 3. Higher frequencies of WT- and alpha-reactive CD8+ T cells among the PASC study group

Next, we analyzed SARS-CoV-2 reactive T cell immunity. As the majority of the study participants were infected with either the WT or the alpha variant, we addressed the WT and alpha reactive CD4+ and CD8+ T cell response.

The frequencies of WT- and alpha-reactive defined as CD4+CD154+CD137+ T cells were similar among the control and PASC study groups (fig. 3A & C). SARS-CoV-2-reactive CD4+ T cells producing IL2, TNFα and GrB showed similar frequencies between the two cohorts (fig. S3A, I, M, C, G, K, O), except for the IFNγ producing WT reactive CD4+ T cells, which showed significantly higher frequencies among the PASC patients (fig. S3E).

**Figure 3:**
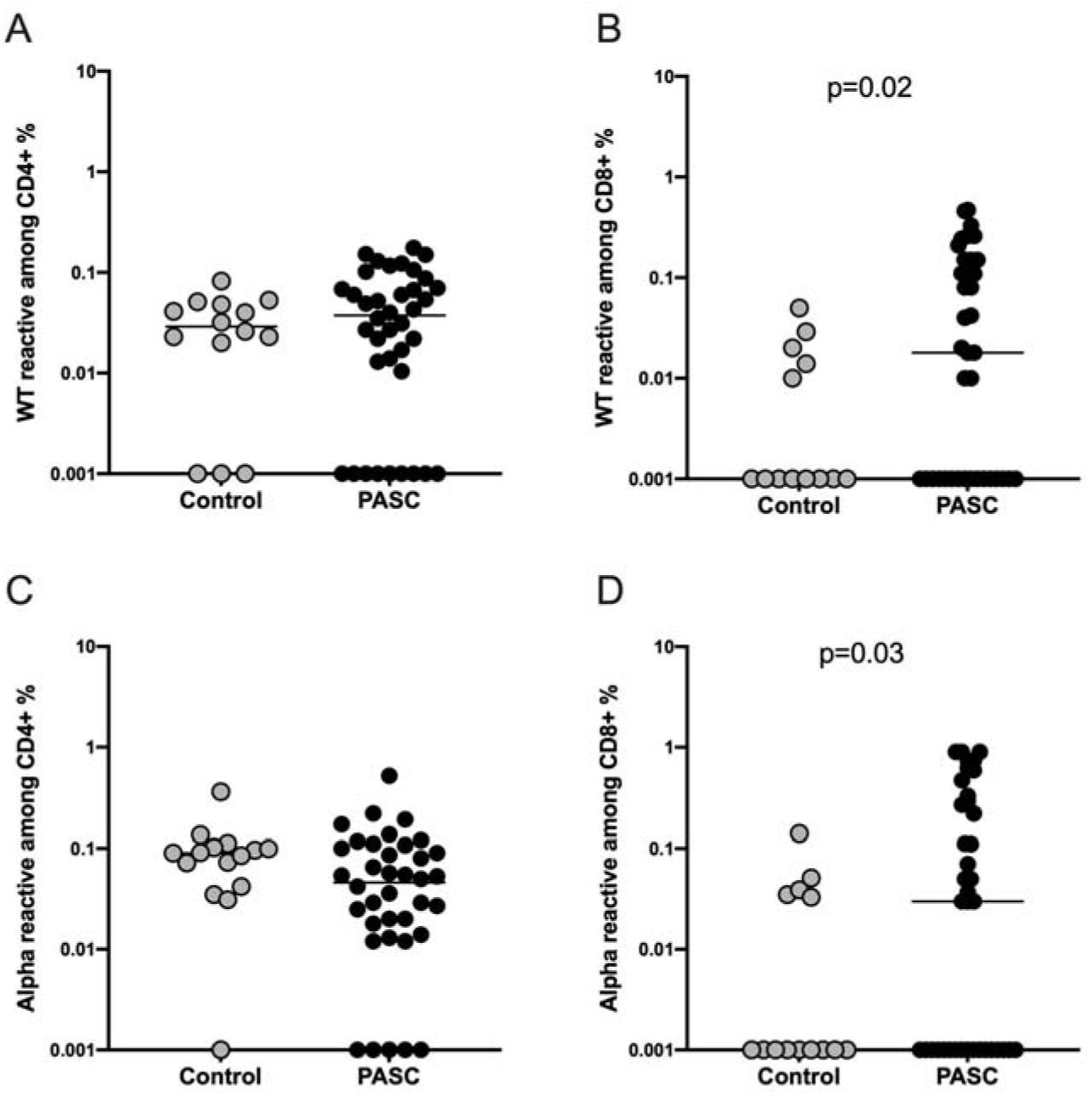
Higher frequencies of SARS-CoV-2 WT and alpha strain reactive CD8+ T cells among the PASC study group. Characterization of SARS-CoV-2 S-reactive T cells in PASC and control subjects. Blood samples of 40 PASC patients and 15 convalescent controls were stimulated with SARS-CoV-2 S-WT and S-Alpha and analyzed by flow cytometry. (A-B) Frequencies of WT- and alpha-reactive CD4+ T cells among PASC and controls. (C-D) Frequencies of WT- and alpha-reactive CD8+ T cells among PASC and controls. SARS-CoV-2 S-reactive CD4+ and CD8+ T cells are defined as CD4+CD154+CD137+ and CD8+CD137+ cells respectively. Antigen-reactive responses were considered positive after the non-reactive background was subtracted, and more than 0.01% were detectable. Scatterplots show line at median. Unpaired data were compared with Mann-Whitney-test. P<0.05 was considered significant, only significant p values are documented in the figures.

On the contrary, the frequencies of WT- and alpha-reactive defined as CD8+CD137+ T cells were significantly higher among the PASC compared to the controls (fig. 3B & D). In the same pattern as above, specific CD8+ T cells producing IL2, TNFα and GrB showed similar frequencies between the two cohorts (fig. S3F, J, N, D, L, P), except for the WT-reactive CD8+ IL2 producing T cells (fig. S3B) and alpha-reactive CD8+ IFNγ producing T cells (fig. S3H), which showed significantly higher frequencies among the PASC patients.

The memory subsets among SARS-CoV-2 reactive T cells, defined by the expression or absence of CD45RA and CCR7, showed no differences among the two cohorts, excluding the WT reactive CD8+ T_EMRA_ cells. The frequencies of WT reactive CD8+ T_EMRA_ cells were significantly higher in PASC (fig. S4).

### 4. Presence of a low avidity CD8 T cell response among the PASC patients

We also performed an analysis of the avidity of SARS-CoV-2-reactive T cells analyzing CD3low (high avidity) and CD3high (low avidity) subsets among reactive CD4+ and CD8+ T cells (gating strategy, fig. S2) as applied before (21, 23). We detected similar frequencies of reactive CD4+ and CD8+CD3low reactive T cells among PASC and controls (fig. 4), indicating that both study groups can achieve similar maximum functional avidity status among the reactive T cell populations. However, the analysis of the CD3high subsets, demonstrated WT- and alpha-reactive CD8+CD3high T cells with significantly higher frequencies among the PASC study group, suggesting the presence of a reactive however with low avidity -potentially uncoordinated-CD8 T cell response among the PASC patients (fig. 5).

**Figure 4:**
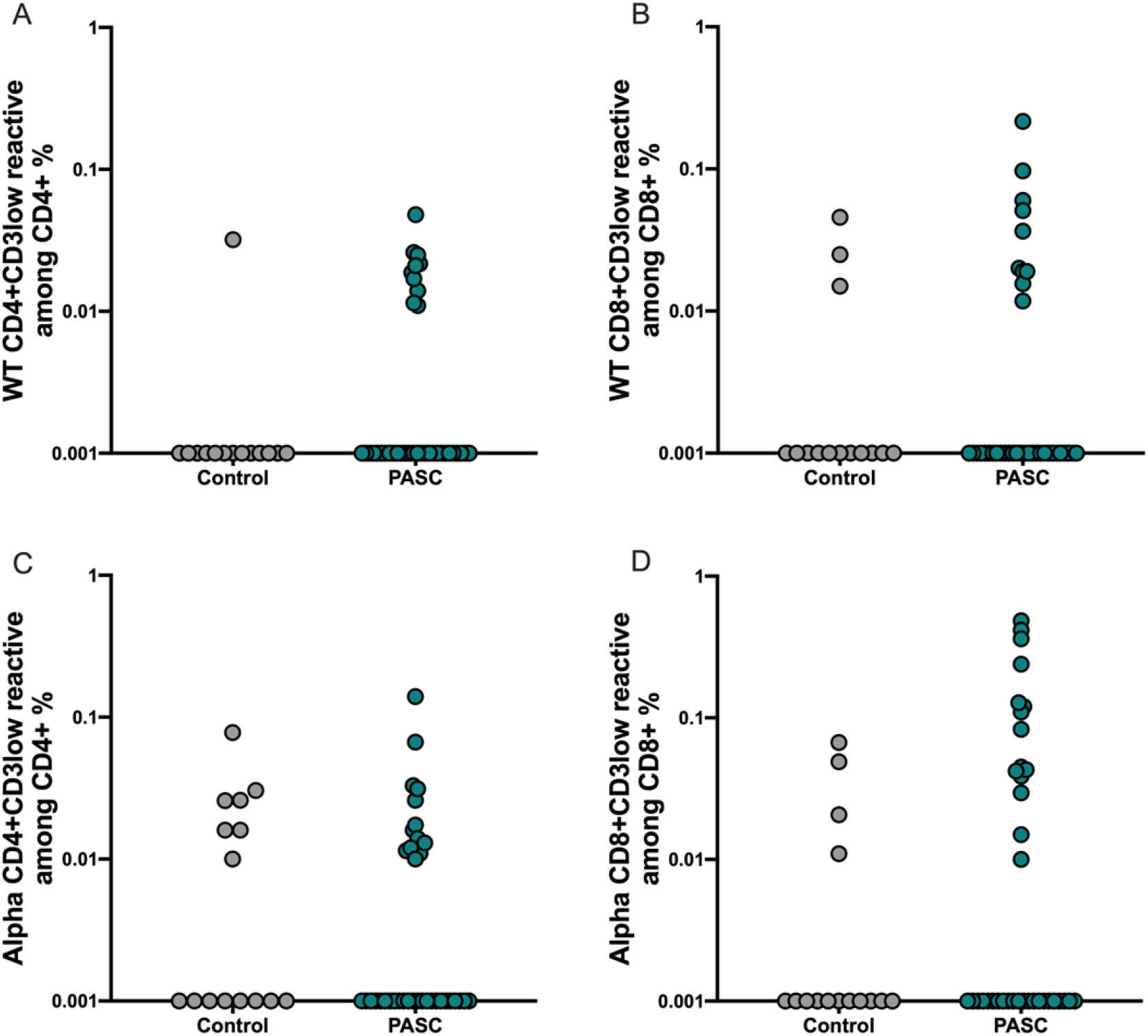
Similar maximal functional avidity among PASC and controls. The maximum functional avidity of SARS-CoV-2 reactive T cells was approached by determining the CD3low+ cells among CD4+CD154+CD137+ and CD8+CD137+ cells. (A-B) Frequencies of WT reactive CD4+CD3low+ and CD8+CD3low+ T cells (C-D) Frequencies of alphareactive CD4+CD3low+ and CD8+CD3low+ T cells.

**Figure 5:**
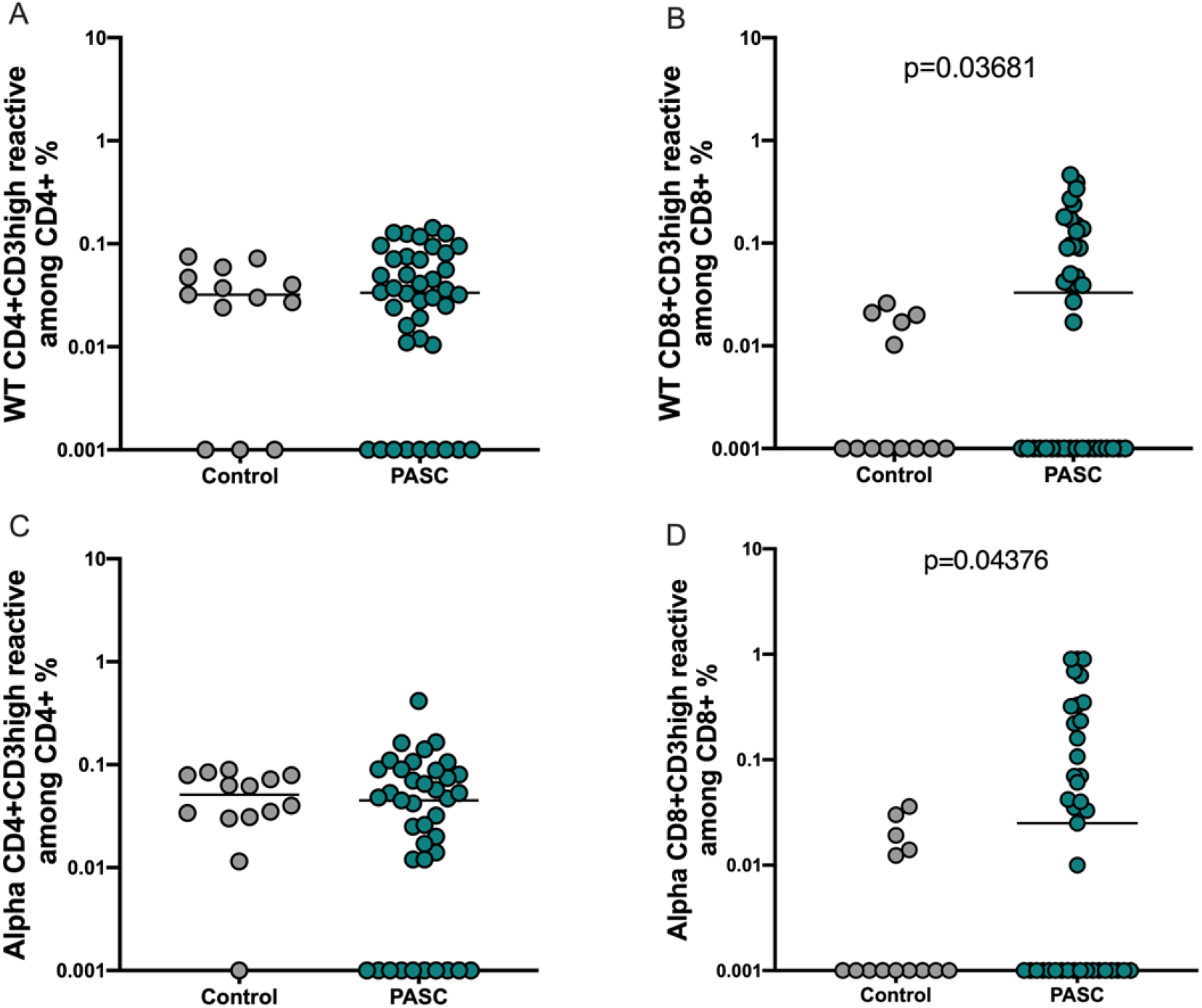
Presence of low avidity CD8 T cell response among the PASC patients. Deteriorated avidity capacity of SAR-CoV-2 reactive T cells was determined by detecting CD3high+ cells among CD4+CD154+CD137+ and CD8+CD137+ cells. (A-B) Frequencies of WT reactive CD4+CD3high+ and CD8+CD3high+ T cells (C-D) Frequencies of alphareactive CD4+CD3high+ and CD8+CD3high+ T cells.

### 5. PASC humoral immunity is not inferior compared to controls

High neutralizing antibodies titers are considered to protect effectively against SARS-CoV-2 infection, and their waning is related with high risk of reinfection or vaccine breakthrough infection (24). Currently it is unclear, whether the NAbs play a role in PASC pathogenesis, therefore we measured the titers of WT and alpha Nabs in PASC and control individuals. We found similar titers of Spike IgGs, WT and alpha NAbs among the two study groups indicating that protective humoral immunity is not impaired among the PSC group (fig. S5).

### 6. High titers of MasR, M5R and PAR2 circulating autoantibodies among the PASC patients

Acute COVID-19 disease has been associated with autoimmunity and the circulation of various types of autoantibodies against nuclear components, phospholipids, coagulation molecules and cytokines (25–28). In the frame of ME/CFS autoantibodies were also demonstrated to have a pathogenic role, and in addition to the above-mentioned molecules, autoantibodies against neuronal components, neurotransmitter and other G-protein coupled receptors have also been described (29–32).

Therefore, we screened the serum of the study participants for circulating IgG autoantibodies against 12 different G protein-coupled receptors; ß2-adr-R, Brady-R1, ATR1, ATR2, MasR, a1-adr-R, CXCR3, ETAR, ETBR, M5R, PAR1, PAR2, two proteins; ACE2, Annexin 2 and the transmembrane receptors Annexin 2R and STAB1. We detected significantly higher titers of MasR, M5R and PAR2 antibodies compared to the controls (fig. 6).

**Figure 6:**
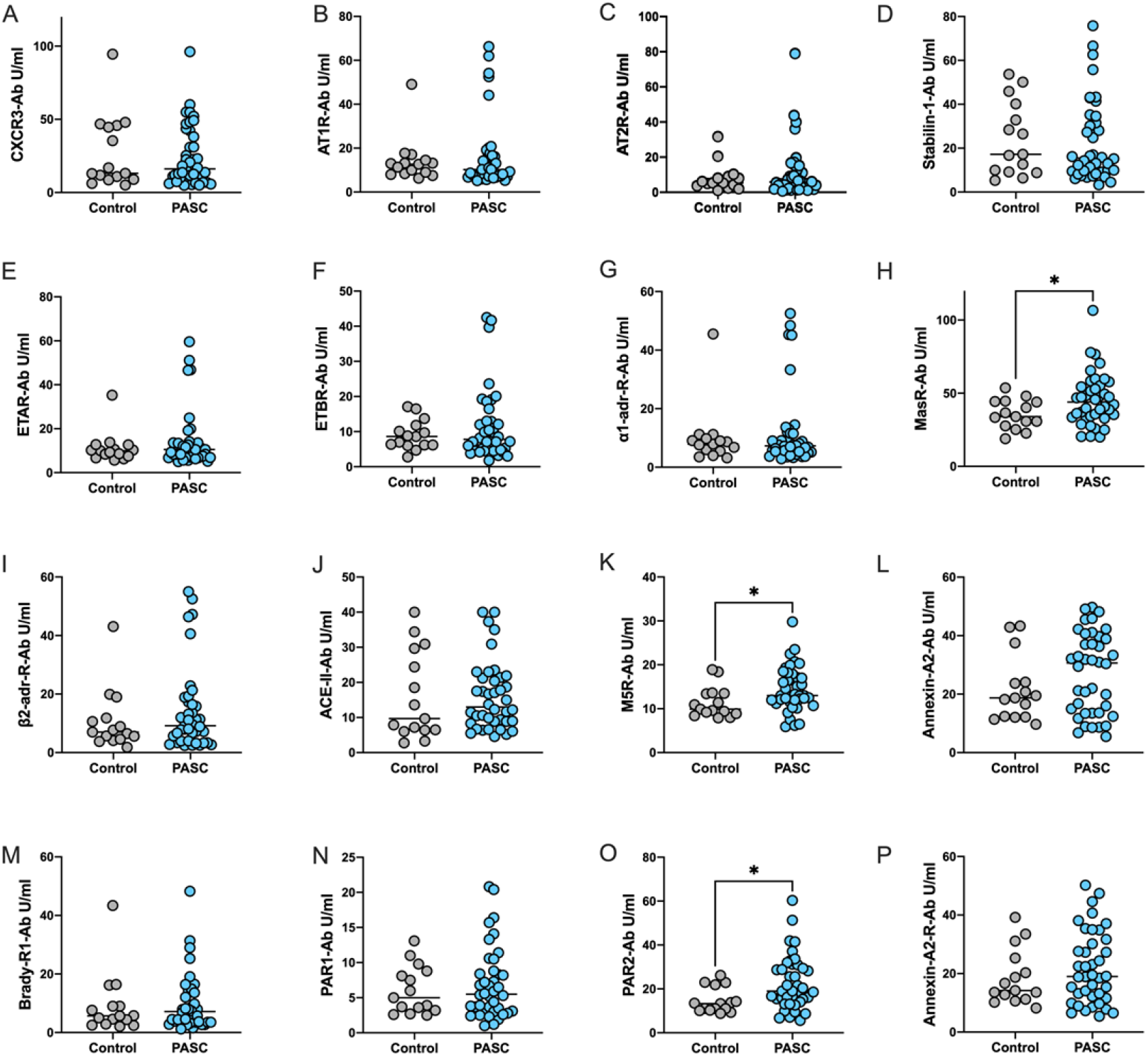
High titers of MasR, M5R and PAR2 circulating autoantibodies among the PASC patients. The serum of the study participants was screened for circulating IgG autoantibodies against 12 different G protein-coupled receptors; ß2-adr-R, Brady-R1, ATR1, ATR2), MasR, a1-adr-R, CXCR3, ETAR, ETBR, M5R, PAR1, PAR2, two proteins; ACE2, Annexin 2 and the transmembrane receptors Annexin 2R and STAB1. Scatterplots show line at median. Unpaired data were compared with Mann-Whitney-test. P<0.05 was considered significant, only significant p values are documented in the figures.

## Discussion

Here, we performed an immune profiling of the cellular and humoral immune response in 40 PASC patients with non-specific PASC. We demonstrate a stronger low avidity SARS-CoV-2 reactive CD8+ T cell response characterized by T_EMRA_ memory phenotype and IFNγ production among the PASC patients compared to controls. The expansion of cytotoxic CD8+ T cells is already demonstrated from independent groups to be an important pathogenic component of gastrointestinal and pulmonary PASC (18,33–34). Regarding the nervous system, it has been proven that upregulation of CD8+ T cells contributes to autoimmunity driven demyelination and axonal damage (35–38), while highly differentiated effector CD8+ TEMRA cells have been associated with neuroinflammation and degeneration (39–40). In agreement with our findings of higher frequencies of IFNγ producing CD8+ T cells among PASC, independent groups also detected higher production of TNFα, IFNγ and IL-6, among PASC patients (34, 41–43).

Furthermore, we analyzed the role of the EBI2, CXCR3 and CCR5 molecules in the frame of PASC. Briefly, CXCR3 is activated by three interferon-inducible ligands CXCL9, CXCL10 and CXCL11 playing an important role in T cell trafficking and function in inflammatory lesions (38, 44–45). EBI2 is expressed with a strong predominance in CD4+ T and B cell memory subsets (46–48) but also in astrocytes (49) and has a predominant role in antigen recognition, proliferative expansion, migration, direct effector activity and regulation of neuronal inflammation (49–50). CCR5 has multiple ligands, including CCL3, CCL4 and CCL5 and plays an important role in inflammatory response to infections. Beside immune cells, CCR5 is highly expressed in microglia and to a lesser extent in astrocytes and neurons(51).

We detected increased frequencies of EBI2+, CXCR3+ and CCR5+ expressing CD4+ and CD8+ T cells among PASC patients indicating their increased activation and migratory capacity. We hypothesize that the upregulation of EBI2+, CXCR3+ and CCR5+ molecules triggers the homing of cytotoxic CD8+ T cells and potentially migration to central nervous system and therefore initiation or persistence of inflammation. In agreement with our findings, Etter et al and Casteneda et al demonstrate elevated pro-inflammatory proteins in cerebrospinal fluid, microglia activation markers and persistent loss of oligodendrocytes and myelinated axons in PASC patients and mice with neurological PASC manifestation(52–53).

Our data are in line with results of other groups demonstrating association of the EBI2 upregulation with neuroinflammation and autoimmunity(49). EBI2 has been shown to be upregulated in a subtype of ME/CFS (54–56), while in multiple sclerosis it has proven to be functionally expressed with a distinct EBI2 expression to lymphocytes migration correlation pattern, also in the absence of Epstein-Barr virus reactivation (57). In the same line, upregulation of the CXCR3 receptor has been associated with neuronal axonal damage secondary to inflammatory response of T cells a) during or after viral infection (37,58) as well as b) in the frame of autoimmunity (59) mainly in multiple sclerosis patients (36,38). Accumulating data indicate the crucial role of CXCR3 in directing the migration of inflammatory mainly CD8+ T-cells into the central nervous system in a CXCR3-dependent manner (45,58,60). Finally, the CCR5 receptor was found to be a negative modulator of learning and memory and its inhibition has been shown to enhance learning, memory, and plasticity processes (61–62). Thus, it was recently shown that CCR5 inhibition with clinically utilized FDA-approved CCR5 antagonist in AIDS therapy (maraviroc) promotes functional recovery in stroke and traumatic brain injury (62–64).

While cellular response directed against SARS-CoV-2 antigens was increased in PASC patients, titers of neutralizing antibodies were not higher in PASC group compared to controls. This suggests that SARS-CoV-2 specific humoral immunity is not involved in the pathogenesis of PASC.

On the other hand, we demonstrate higher titers of MasR, M5R and PAR2 circulating autoantibodies among the PASC patients. GPCR antibodies are dysregulated in various autoimmune and non-autoimmune diseases. Their upregulation may indicate alterations in receptor expression or altered function. By acting as ligands to their target receptors, antibodies against GPCRs can modulate receptor signaling and were mostly shown to result in agonist stimulation, but their function was also found to be impaired in ME/CFS(65–66). A recent study showed that several GPCR antibodies are elevated in acute COVID-19 and CXCR3 and angiotensin1 receptor AGTR1 antibodies to have the strongest association with disease severity (25). Elevated muscarinic acetylcholine receptor antibodies are demonstrated in a subset of ME/CFS patients (29–30, 67–68). M5R is an important vasodilator in the brain (69–70), and elevated levels may indicate vascular dysregulation, however further studies are needed to assess its function in PASC. An impaired circulation and oxygen supply could result in many symptoms of ME/CFS and PASC (71–73). We also demonstrate significantly higher titers of MasR antibodies. ACE2/Ang-(1–7)/MasR axis activation inhibits inflammation and fibrosis and counterplays the ACE/angiotensin II/AGTR1/RAS axis leading to vasodilatation (74–77). The upregulation of ACE/angiotensin II/AGTR1/RAS axis with concurrent reduction of ACE2/Ang-(1–7)/MasR axis has been associated with unfavorable COVID-19 outcome during acute infection (25). Similar to elevated M5R, MasR antibodies may indicate vascular dysregulation. Lastly, PAR2 is expressed on several immune cells and its role is double-edged: its upregulation on the one hand triggers neurogenic inflammation, hyperalgesia and metabolic dysfunction, whereas it leads to protective effects in gastrointestinal and pulmonary mucosa (78–82). We suggest that the PAR2 antibodies function agonistically in the frame of PASC, however further studies are needed to assess our indirect results. Interestingly, Sotzny et al correlate levels of F2R/PAR-1 antibodies with the severity of cognitive symptoms and of M5R with fatigue in post COVID ME/CFS (83).

There are some limitations of this study that should be addressed. The most important limitation is the slightly unbalanced presentation of some demographic characteristics. Firstly, the PASC study group was significantly older compared to controls. As aging is related to immunosenescence and chronic inflammation (84–85) we cannot completely exclude that our results may reflect the phenomenon of inflammaging in PASC. Additionally, the PASC study group presented more metabolic comorbidities compared to controls. Moreover, BMI among PASC study cohort was significantly higher compared to controls and might, therefore, also bias our observation. Obesity has been correlated with increased risk for PASC development (86–87) and is linked to adipose chronic tissue inflammation characterized by infiltration and activation of immune cells that overproduce cytokines and chemokines (88–92). Lastly, future studies exploring correlational relationships of T cell response with the clinical severity of non-specific PASC by implementing neurological assessment tools, would be of interest.

According to Taquet et al, non-specific PASC symptomatic shows no clinical excess when compared to other respiratory infections (93). However, its occurrence is more frequent, as COVID-19 incidence is higher compared to other respiratory infections (94). Beside the burden of PASC on the health care system, non-specific PASC has been correlated to reduced working hours and inability to work in 22% of PASC patients (95). Despite its benignity in the majority of the cases, non-specific PASC is a public health and social issue requiring interdisciplinary attention. In summary, our data suggest that the origins of non-specific PASC may derive from an inflammatory response triggered by a strong however low avidity SARS-CoV-2 reactive pro-inflammatory CD8+ T cell response, and generation of circulating autoantibodies. Our findings may have implications on future therapeutical strategies and public health policies.

## Study Approval

The study was approved by the Ethics Committee of University Hospital Essen (20-9753-BO). Written informed consent was obtained from all participants.

## Author Contributions

KP, NB, MA and US participated in research design. KP, MK, MZ, HR, SD and AG participated in data curation and sample acquisition. KP, CS and NB participated in the writing of the paper. NB, OW, TW, US and HD participated in funding acquisition and project administration. KP, MA, TLM, SP, HH, JJ and AK participated in the performance of the research. NB, TW, OW and US contributed new reagents or analytic tools. KP, NB and MA participated in data analysis. All authors contributed to the article and approved the submitted version.

## Data Availability

The datasets generated during and/or analyzed during the current study are available from the corresponding author on reasonable request.

## Supplemental Table of Contents

### Supplementary Tables

**Table S1:**
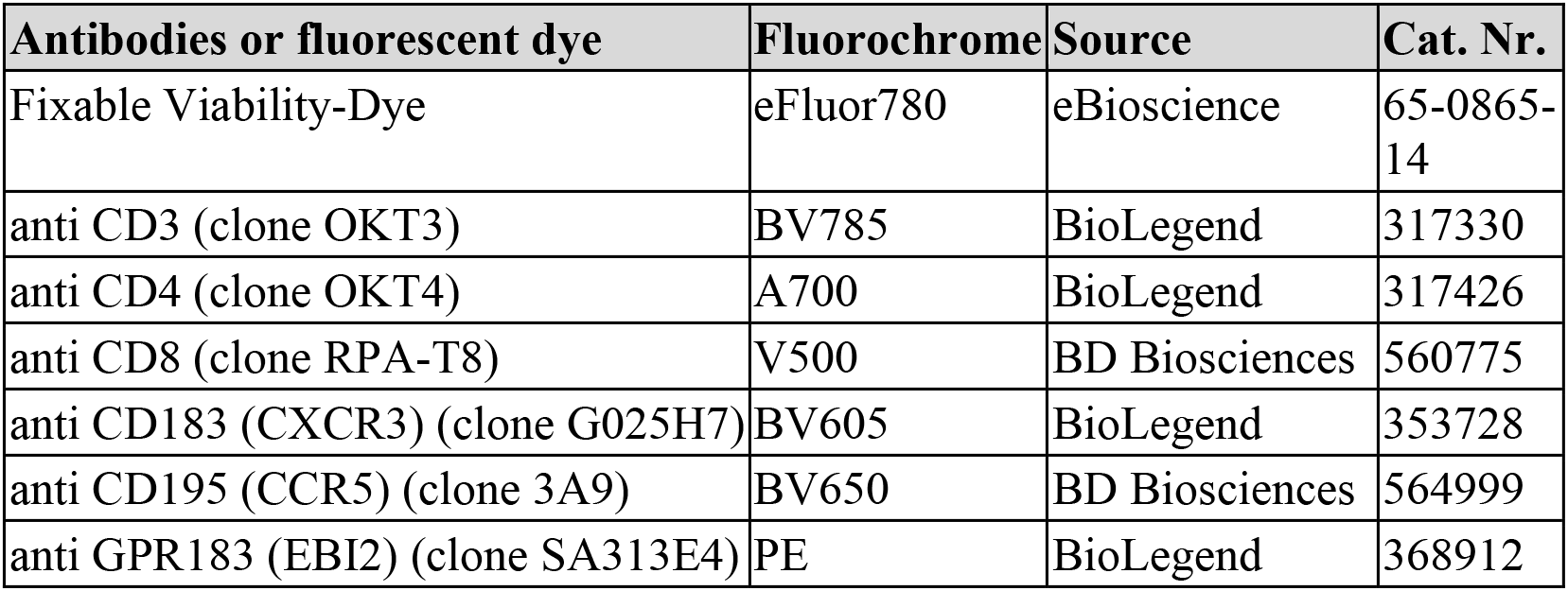
Fluorochrome coupled antibodies and fluorescent dye for analysis of EBI2+, CCR5+ and CXCR3+ T cells

| Antibodies or fluorescent dye | Fluorochrome | Source | Cat. Nr. |
| --- | --- | --- | --- |
| Fixable Viability-Dye | eFluor780 | eBioscience | 65-0865-14 |
| anti CD3 (clone OKT3) | BV785 | BioLegend | 317330 |
| anti CD4 (clone OKT4) | A700 | BioLegend | 317426 |
| anti CD8 (clone RPA-T8) | V500 | BD Biosciences | 560775 |
| anti CD183 (CXCR3) (clone G025H7) | BV605 | BioLegend | 353728 |
| anti CD195 (CCR5) (clone 3A9) | BV650 | BD Biosciences | 564999 |
| anti GPR183 (EBI2) (clone SA313E4) | PE | BioLegend | 368912 |

**Table S2:** Fluorochrome coupled antibodies and fluorescent dye for analysis of SARS-CoV-2 reactive T cells

| Antibodies or fluorescent dye | Fluorochrome | Source | Cat. Nr. |
| --- | --- | --- | --- |
| Fixable Viability-Dye | eFluor780 | eBioscience | 65-0865-14 |
| anti CCR7 (clone G043H7) | PerCP-Cy5.5 | BioLegend | 353220 |
| anti CD4 (clone OKT4) | A700 | BioLegend | 317426 |
| anti CD8 (clone RPA-T8) | V500 | BD Biosciences | 560775 |
| anti CD45RA (clone HI100) | BV605 | BioLegend | 304134 |
| anti Granzyme B (clone GB11) | FITC | BioLegend | 515403 |
| anti IL2 (clone MQ1-17H12) | PE | BioLegend | 500307 |
| anti CD185(CXCR5) (clone MP4-25D2) | PE-Dazzle594 | BioLegend | 356927 |
| anti CD137 (4-1BB) (clone 4B4-1) | PE-Cy7 | BioLegend | 309818 |
| anti CD154 (CD40L) (clone 24-31) | A647 | BioLegend | 310818 |
| anti TNFα (clone MAb11) | eFluor450 | eBioscience | 48-7349-42 |
| anti IFNγ (clone 4S.B3) | BV650 | BioLegend | 502538 |
| anti CD3 (clone OKT3) | BV785 | BioLegend | 317330 |

### Supplementary Figures

**Figure S1:**
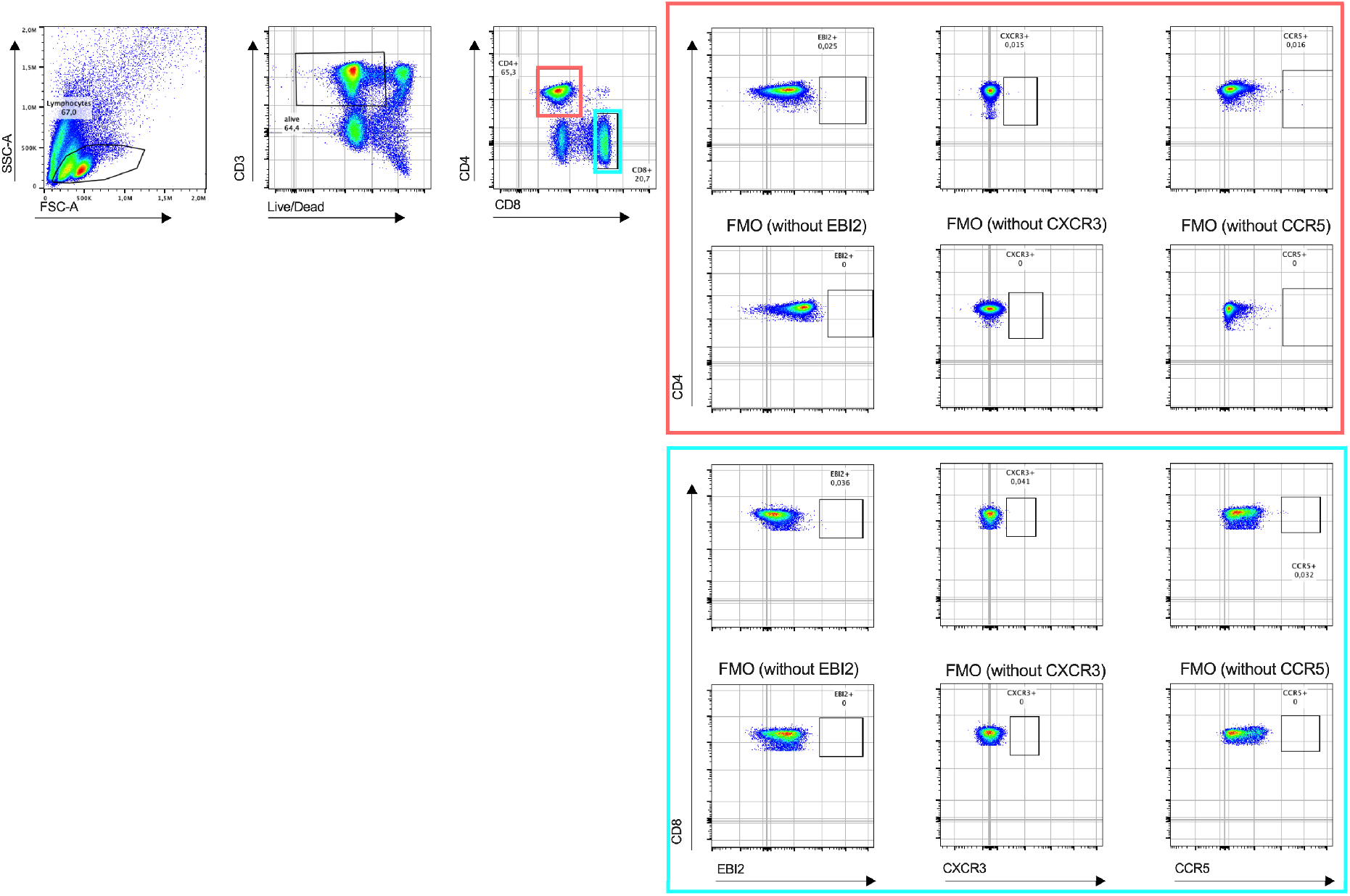
Flow cytometry gating strategy for identification of activated/migratory T cells. Thawed and rested overnight PBMCs were stained with optimal concentrations of antibodies. Living single lymphocytes were analyzed for expression of CD3, CD4, and CD8. The expression of the activation and migration markers EBI2, CCR5 and CXCR3 was assessed on CD4+ (orange boxes) and CD8+ (blue boxes) single positive T cells. Fluorescence minus one controls (FMO) were used for optimal gating of EBI2+, CCR5+ and CXCR3+ populations. Representative example of 40 patients with post COVID-19 syndrome and 15 healthy convalescent individuals is shown. Plots of a PASC study subject are depicted.

**Figure S2.**
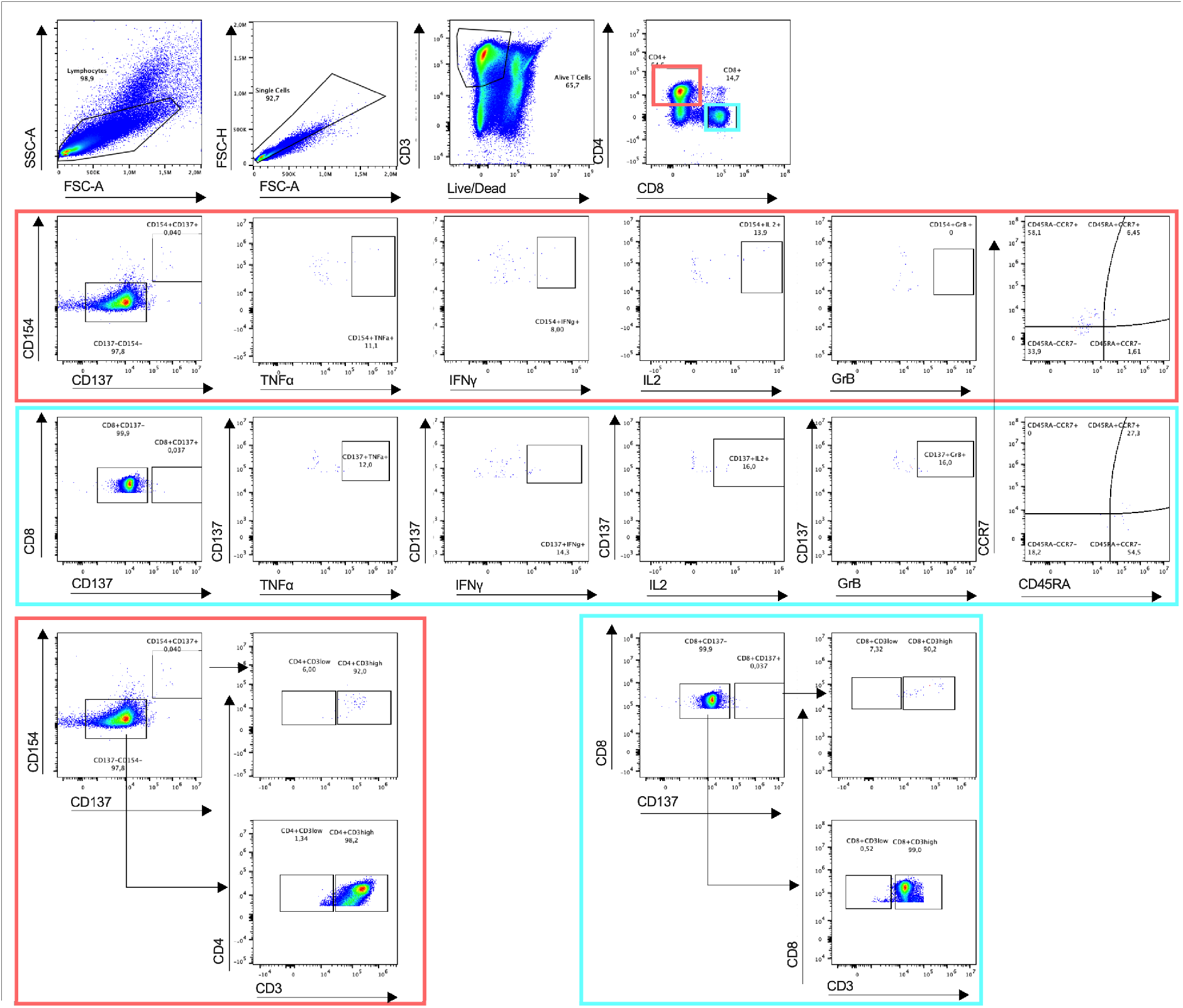
Flow cytometry gating strategy for identification and quantification of SARS-CoV-2 reactive T cells. PBMCs were stimulated for 16 h with one of the following SARS-CoV-2 peptides spanning *in silico* predicted immunodominant parts of the; WT Spike SARS-CoV-2 and the B.1.1.7 D614G Spike mutant. After 2 h, Brefeldin A was added to the culture to block secretion of cytokines and effector molecules. Living single lymphocytes were analyzed for expression of CD3, CD4, and CD8. CD4+ T cells (orange boxes) were analyzed for the expression of CD154 and CD137. CD8+ T cells (blue boxes) were analyzed for expression of CD137. Both CD4+ and CD8+ T cells were further analyzed for the production of cytokines IFNγ, TNFα, IL2 and GrB. Evaluation of the memory subsets was performed using the markers CCR7 and CD45RA (T_CM_=CD45RA-CCR7+, T_NAIVE_=CD45RA+CCR7+, T_EM_=CD45RA-CCR7-T_EMRA_=CD45RA+CCR7-). Furthermore, CD4+CD154+CD137+, CD8+CD137+ and cells were analyzed for the expression of CD3low and CD3high. Representative example of 40 patients with post COVID-19 syndrome and 15 healthy convalescent individuals. Plots of a PASC study subject are depicted.

**Figure S3:**
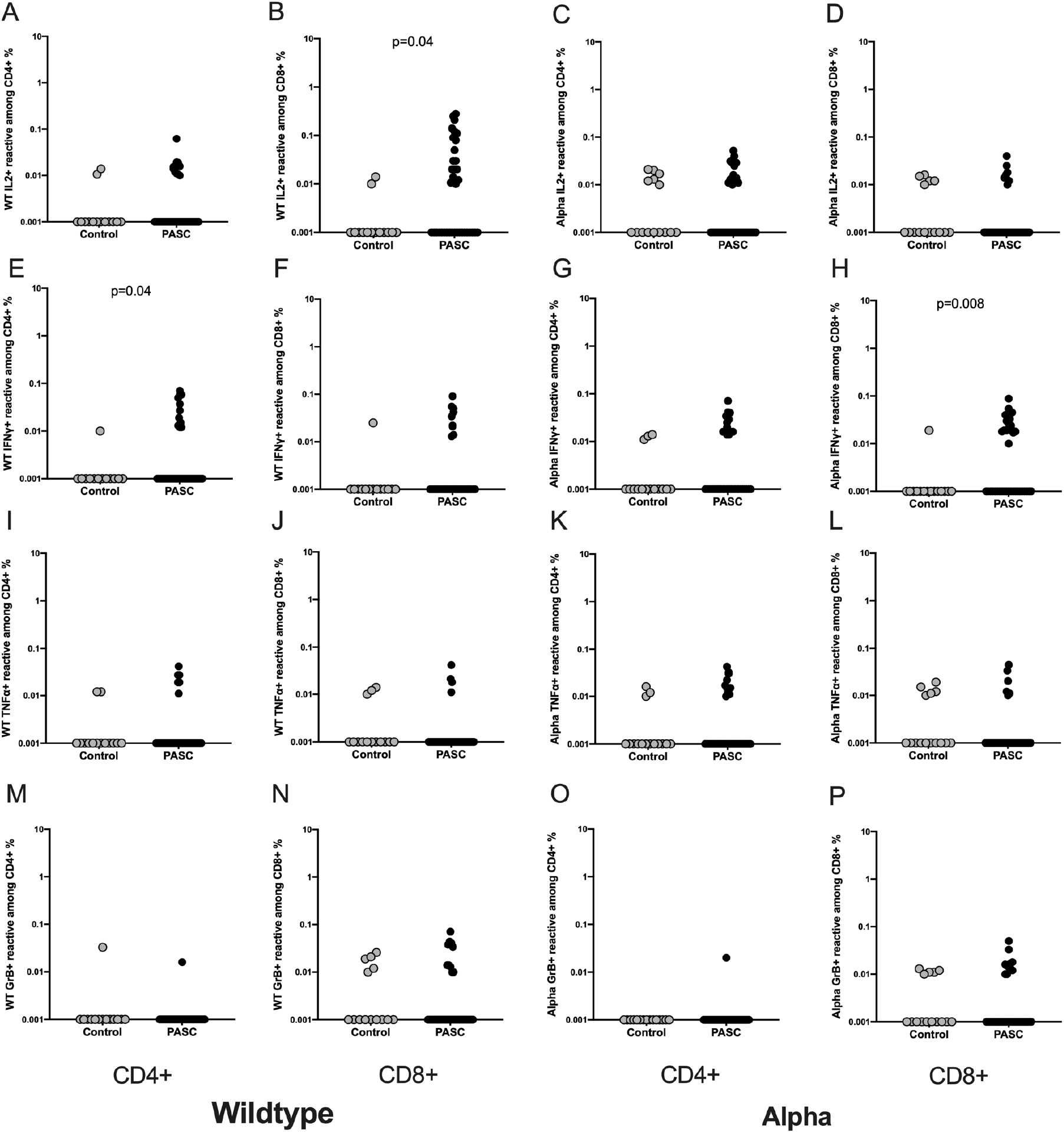
Higher frequencies of IFNγ producing CD4+ and CD8+ reactive T cells. The frequencies of IL2, IFNγ, TNFα or GrB producing WT- and alpha-reactive T cells were analyzed among PASC subjects and controls. (A-D) IL2 producing SARS-CoV-2 reactive CD4+ and CD8+ T cells. (E-H) IFNγ producing SARS-CoV-2 reactive CD4+ and CD8+ T cells. (I-L) TNFα producing SARS-CoV-2 reactive CD4+ and CD8+ T cells. (M-P) GrB producing SARS-CoV-2 reactive CD4+ and CD8+ T cells.

**Figure S4:**
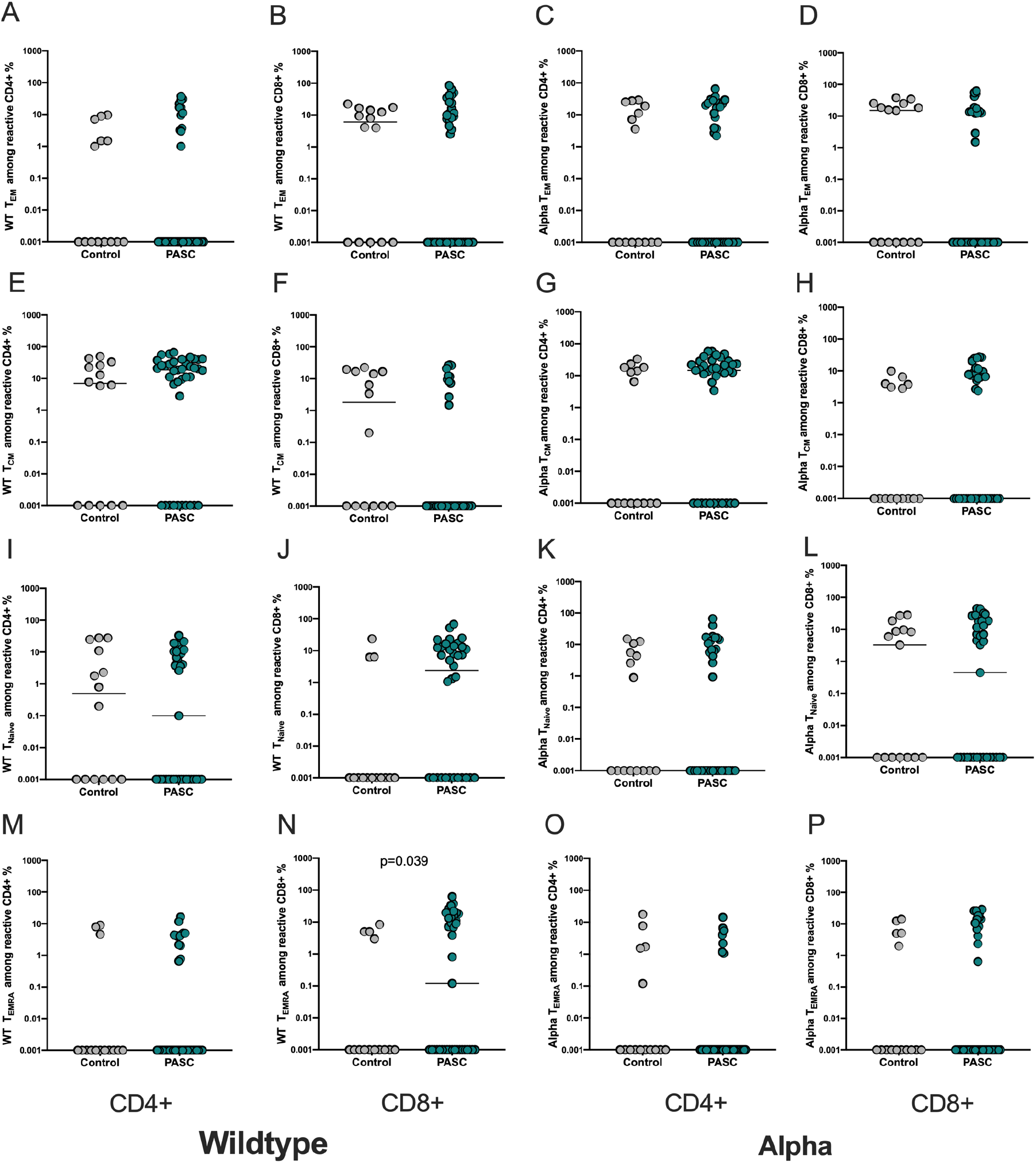
Higher frequencies of WT-reactive CD8+ T_EMRA_ cells among the PASC study group. Evaluation of the memory subsets was performed using the markers CCR7 and CD45RA (T_CM_=CD45RA-CCR7+, T_NAIVE_=CD45RA+CCR7+, T_EM_=CD45RA-CCR7-T_EMRA_=CD45RA+CCR7-). Frequencies of the memory subsets of WT- and alpha-reactive CD4+ and CD8+ T cells in PASC subjects were compared to controls. (A-D) WT- and alpha-reactive CD4+ and CD8+ T_EM_ cells. (E-H) WT- and alpha-reactive CD4+ and CD8+ T_CM_ cells. (I-L) WT- and alpha-reactive CD4+ and CD8+ T_NAIVE_ cells. (M-P) WT- and alpha-reactive CD4+ and CD8+ T_EM_ cells.

**Figure S5:**
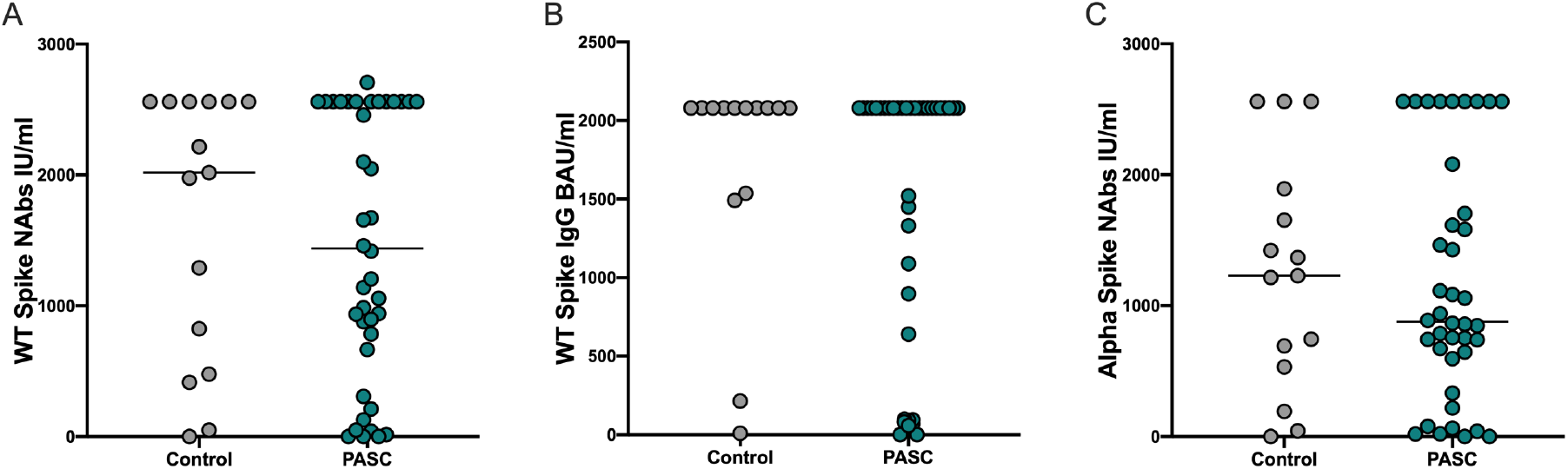
PASC humoral immunity is not inferior compared to controls. Analysis of WT Spike IgG and WT and Alpha NAbs titers of both study groups. (A) Spike IgG titers. (B) WT NAbs titers. (C) Alpha NAbs titers. Scatterplots show line at median. Unpaired data were compared with Mann-Whitney-test. P<0.05 was considered significant, only significant p values are documented in the figures.

